# Non-targeted Analysis of Extracellular Vesicle-Enriched Plasma Proteome between Early and Late Rugby Playing Career

**DOI:** 10.1101/2025.03.10.642401

**Authors:** Abhishek Jagan, Yusuke Nishimura, Tim Donnovan, Jatin G Burniston

## Abstract

Rugby players may repeatedly incur high-impact collisions that could predispose them to neurodegenerative conditions but the processes underlying the heightened risk are currently unclear. This project investigates whether the proteome of plasma extracellular vesicles (EV) could carry putative diagnostic biomarkers to indicate differences in risk to neurodegenerative conditions across a rugby playing career.

Twenty-four males were recruited, including eight academy players (18 ± 1 y), eight professional rugby players (33 ± 5 y) with >10-year rugby career and eight CrossFit athletes (32 ± 5 y) with no history of collision-related sports injuries. Membrane-bound particles (i.e. EV) were enriched from plasma using hyper-porous strong-anion exchange magnetic microparticles and tryptic peptides were analysed using nano-flow liquid chromatography and high-resolution tandem mass spectrometry. Differences in protein abundance were investigated by one-way analysis of variance (with correction for multiple testing) after label free quantitation.

In total, 449 proteins were confidently identified (false discovery rate; FDR <1 %) and gene ontology profiling confirmed 414 of these proteins were of EV origin. One-way ANOVA highlighted 128 significantly (P<0.05, q<0.02) different proteins across the three participant groups, of which 31 proteins were specific to professional rugby players. Seven of these proteins (APOA1, APOM, CLUS, BIP, VCAM1, NID1 and MMP9) which were depleted and one protein ZPI which was elevated have previously recognised roles in neurodegenerative processes.

In conclusion, non-targeted analysis highlighted that proteins associated with neuroprotection were specifically depleted in the plasma EV proteome of long-serving professional rugby players compared to younger academy rugby players or age-matched cross-fit athletes that did not have a history of collision-related sports injuries. Our findings shed new light on processes affected by a professional rugby playing career, further application of this type of analysis could be used to develop biomarker panels useful for predicting at-risk athletes or for guiding treatment interventions.

## Introduction

Rugby has a high injury incidence rate compared to other team contact sports and players are regularly exposed to collision injuries, including blunt trauma to the upper body, head and neck. Sport-related concussion (SRC) is the most common (20 % of all match injuries) injury in Rugby Football Union in England (McCrory et al., 2017) and SRC incidence rates are rigorously monitored. The England professional Rugby injury surveillance project (PRISP, 2024) reported 18.4 incidences of SRC per 1000 hours of professional play during the 2022-23 season. Reliable diagnosis of concussion is the first step in the management of player health but diagnosis can be challenging in the context of team contact sports. The diagnostic criteria for concussion include actual or suspected loss of consciousness (Broglio et al., 2022), which may not always occur and players may also present co-occurring collision or crush injuries to the torso or neck.

The long-term consequences of SRC remain to be fully understood but connections have been drawn between concussions and negative consequences on cognitive function, including chronic traumatic encephalopathy (CTE) (Blecher et al., 2019), Alzheimer’s disease (AD), and motor neuron disease (MND) (Chen et al., 2022). Former elite rugby players may have lower cognitive function compared to non-contact sport players (Hume et al., 2017) and older adults with a history of three or more concussions during adulthood have worse cognitive performance than those with no history of concussion (Gallo et al., 2022).

Electroencephalography (EEG) (Gosselin et al., 2012), functional magnetic resonance imaging (fMRI) (Chen et al., 2008), and advanced magnetic resonance imaging (MRI) (Bartnik-Olson et al., 2014) can each distinguish distinct patterns in individuals with persistent post-concussion symptoms, and there is growing interest in identifying blood biomarkers for SRC diagnosis.

Plasma biomarker studies in rugby players have targeted biomarkers of neurodegeneration such as Tau, neurofilament light (NFL) (Shahim et al., 2018), glial fibrillar acidic proteins (GFAP) (Laverse et al., 2020) and others (Oris et al., 2023), as potential indicators of traumatic brain injuries. These hypothesis-led studies suggest a connection between changes in plasma proteins and the occurrence and recovery of SRC. However, biomarkers, such as S100 Calcium binding protein B (S100B), that are elevated after SRC (Meier et al., 2017), are also known to be elevated by physical activities such as running (Stocchero et al., 2014) and may, therefore, also indicate exercise-induced muscle damage rather than collision-specific processes. Plasma proteomics offers broader insights into the physiological responses of athletes (Zhao et al., 2020; Mi et al., 2023) and there is growing interest in using omic techniques to discover protein biomarkers related to tissue inflammation and cerebrovascular integrity, particularly in athletes with a history of concussions (Meier et al., 2022; Rath et al., 2022).

Extracellular vesicles (EV), including microvesicles and exosomes, have gained attention for their roles in exercise-induced cytokine secretion (Frühbeis et al., 2015) and as mediators of intercellular communication. EV can be characterized by size, cargo, and surface markers and can contain cargoes, including proteins, lipids, metabolites, messenger RNA, micro RNA (miRNA) and nucleic acids (Zaborowski et al., 2015). The unique properties of EV, including their resistance to enzymatic degradation, ability to cross the blood-brain barrier, and traceability to their cell of origin, make them a promising source for traumatic brain injury biomarkers (Banks et al., 2020; Osteikoetxea et al., 2015). Indeed, EV miRNA biomarkers can distinguish between injured and control samples (Ko et al., 2018) and recent EV proteomic studies have identified proteomic signatures in National Football League (NFL) players at risk of CTE (Muraoka et al., 2021).

EV represent a small portion of the plasma proteome and methods for isolating EV, including size exclusion chromatography (Brahmer et al., 2019) and ultracentrifugation (Holcar et al., 2020) are labour-intensive and require specialized equipment. Herein, we employed a magnetic polymeric microparticles (Mag-Net) designed for the isolation, purification, and enrichment of EV from plasma (Wu et al., 2023). EV proteomes were studied across three independent groups of athletes, including: young academy rugby players (Acd), older professional rugby players (Pro) with >10 years experience, and CrossFit (Xfit) athletes age-matched to the Pro players. The Xfit athletes reported no history of impact or concussion injuries and the rugby players had no reported incidence of concussion during the six months prior to the collection of samples for this study.

## Methods

### Participants

Figure 1 A & B provide an overview of the experimental design and analysis protocol. Eight academy players (Acd; 18±1 years) in their third week of preseason and had no reported incidence of concussion from the previous six months, eight Professional rugby players with 10+ years professional rugby experience (Pro; 33±5 years) in the first week of preseason with no concussions reported for the last six months and eight CrossFit athletes (Xfit; 32±5 years) age matched to the Pro group and no history of concussion or other high grade impact injuries, were recruited for blood sampling (Figure 1). All participants were male and gave their informed consent to the procedures approved (M23_SPS_3182) by the Research Ethics Committee of Liverpool John Moores University.

**Figure 1.**
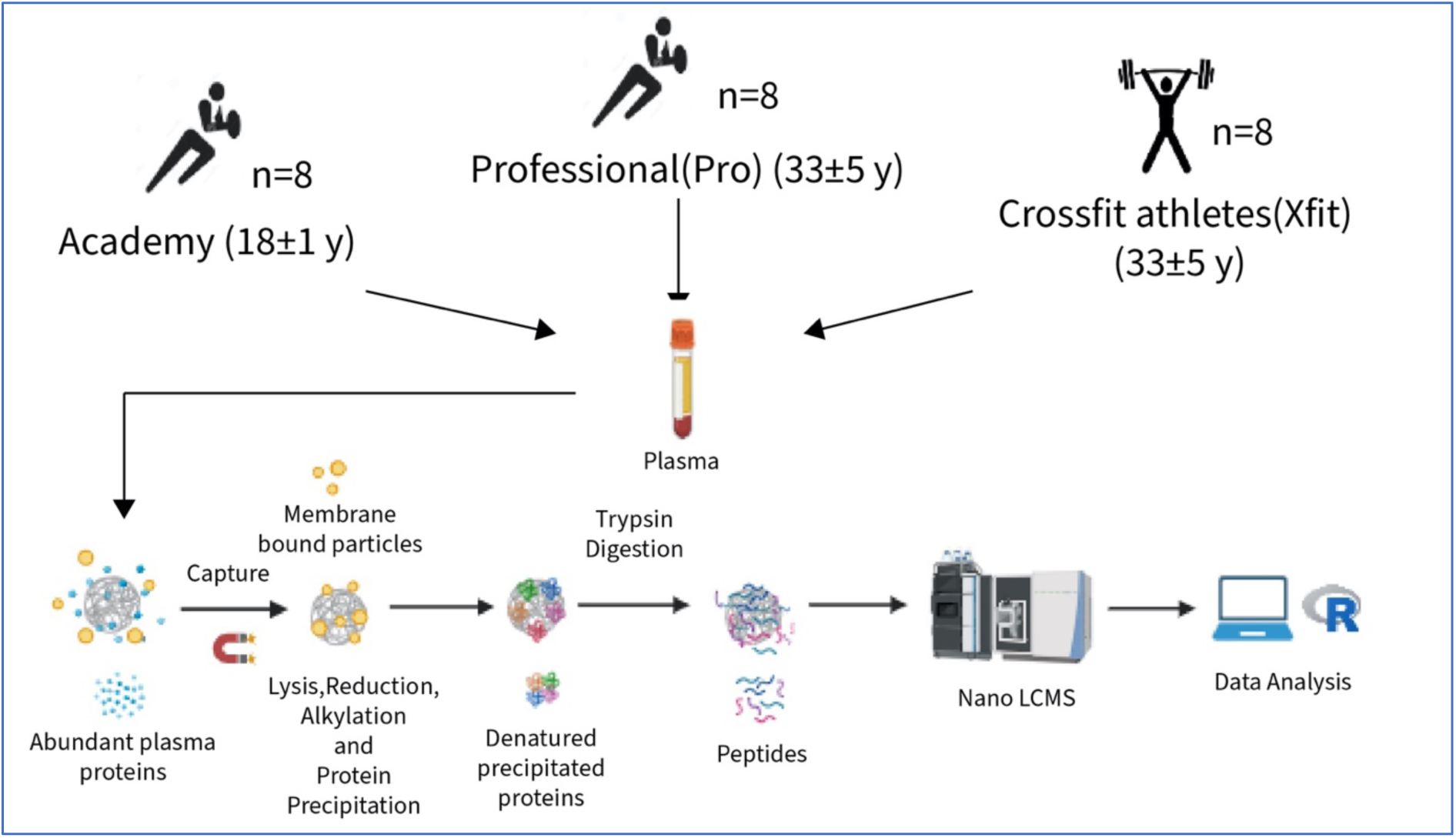
Experimental design and sample processing. Workflow of the quantitative proteomic analysis of the plasma EV isolated using the MagResyn® SAX protocol from healthy male Professional (Pro) players (n=8) with 10+ years of professional rugby playing experience, academy (n=8) players in the 3^rd^ year of their youth development program and CrossFit athletes (Xfit) (n=8) age-matched to professional rugby players with no reported history of concussion or major trauma.

### Blood Samples Collection

Blood samples were collected in vacutainers containing Ethylenediaminetetraacetic acid (EDTA). Blood samples were centrifuged at 1200 × g for 10 mins at 4°C. Plasma was aliquoted and stored at −80°C until further analysis. Freeze/thaw cycles were prohibited.

### Membrane Particle Enrichment

Magnetic affinity-based isolation was used to enrich the membrane particle fraction using a bead based protocol developed by Wu et al (Wu et al., 2023). Twenty-five µL MagReSyn® SAX (Resyn Biosciences, Ranburg, South Africa) (MAG) beads was equilibrated with wash buffer (Wb; 50mM Bis Tris Propane pH 6.3, 150mM NaCl) then mixed with 100 µg plasma protein diluted in binding buffer (Bb; 100 mM Bis Tris Propane pH 6.3, 150 mM NaCl) supplemented with cOmplete™ mini-EDTA free protease inhibitor (Roche). During each incubation, the protein and bead slurry was mixed at 550 rpm (ThermoMixer® Eppendorf, UK/Ireland). After an initial 30-min incubation, beads were washed thrice with 500 µL of Wb then incubated in Lysis and Reduction Mix (LR) (50 mM Tris, pH 8.5, 1% SDS, 10 mM TCEP) for 60 min at 37°C with mixing. Iodoacetamide (15 mM final concentration) was added, and samples were incubated protected from light for 30 mins with mixing. Beads were washed 3 times in acetonitrile (ACN) and 3 times in absolute ethanol prior to on-bead digestion in 25 mM ammonium bicarbonate and trypsin (1:50 enzyme to protein ratio) at 37°C for 2 hours with mixing. Digestion was halted by adding trifluoroacetic acid (TFA) to a final concentration of 0.5 % (v/v). The concentration of the supernatant was estimated using spectrophotometry (260 nm/ 280 nm ratio; N50 Nanophotometer, IMPLEN, Munich, Germany). Peptide aliquots (5 μg) were purified on C_18_ Zip-tips (Millipore) and eluted using a 40:60 mix of ACN and 0.1% TFA. Peptide solutions were dried by vacuum centrifugation (Thermo Scientific, SpeedVac SPD1030) for 40 minutes at 50°C, then reconstituted in 0.1 % formic acid for LC-MS/MS analysis.

### Liquid chromatography-mass spectrometry analysis

Peptide mixtures were analysed using an Ultimate 3000 RSLC nano liquid chromatography system (Thermo Scientific) coupled to Q-Exactive orbitrap mass spectrometer (Thermo Scientific). Samples (700 ng) were loaded on to the trapping column (Thermo Scientific, PepMap 100, 5 μm C18, 300 μm X 5 mm), using ulPickUp injection, for 1 minute at a flow rate of 25 μl/min with 0.1 % (v/v) TFA and 2% (v/v) ACN. Samples were resolved on a 500 mm analytical column (Easy-Spray C18 75 μm, 2 μm column) using a gradient of 97.5 % A (0.1 % formic acid) 5 % B (79.9 % ACN, 20 % water, 0.1 % formic acid) to 70 % A: 30 % B over 120 min at a flow rate of 300 nl/min. The data-dependent selection of the top10 precursors selected from a mass range of m/z 300-1600 was used for data acquisition, which consisted of a 70,000-resolution full-scan MS scan at m/z 200 (AGC set to 1e6 ions with a maximum fill time of 250ms). MS/MS data were acquired using quadrupole ion selection with a 3.0 m/z window, HCD fragmentation with a normalized collision energy of 30 and in the orbitrap analyser at 17,500-resolution at m/z 250 (AGC target 5e4 ion with a maximum fill time of 250ms). To avoid repeated selection of peptides for MS/MS, the program used a 20s dynamic exclusion window.

### Label-Free Quantitation of Protein Abundances

Progenesis Quantitative Informatics for Proteomics (QI-P; Nonlinear Dynamics, Waters Corp., Newcastle, UK) was used for label-free quantitation, consistent with previous studies (Hesketh et al., 2020; Burniston et al., 2014). Log-transformed MS data was normalized by inter-sample abundance ratio, and relative protein abundances were calculated using nonconflicting peptides only. MS/MS spectra were exported in Mascot generic format and searched against the Swiss-Prot database (2023_08) restricted to ‘Homo Sapiens (human)’ (20,424 sequences) using locally implemented Mascot server (v.2.8; www.matrixscience.com). The enzyme specificity was trypsin with 2 allowed missed cleavages, carbamidomethylation of cysteine (fixed modification) and oxidation of methionine (variable modification), m/z error tolerances of 10 ppm for peptide ions and 20 ppm for fragment ion spectra were used. False discovery rate (FDR) was calculated using a decoy database search and only high confidence peptide identifications (>1 % FDR) were retained. The Mascot output (xml format) was recombined with MS profile data in Progenesis.

### Statistical Analysis

All statistical analysis was performed in R version 4.2.1. One-way analysis of variance was performed to assess protein abundance differences across the independent groups. Statistical significance was set at P < 0.05, q-values were used to calculate FDR and account for multiple testing. Fuzzy c-means clustering (Mfuzz package in R) was used to segregate patterns of abundance difference amongst statistically significant proteins. This analysis revealed five distinct cluster patterns, with each protein assigned a membership value (MV) based on its abundance pattern across the three groups.

Bioinformatic Analysis.

Gene ontology (GO) and KEGG pathway analyses were performed against whole human genome using ShinyGO (v0.80) (Ge et al., 2019). Enrichment of GO terms were accepted at an FDR cut-off 0.005 and pathways were restricted to 10 and a pathway size range of 2 to 2000. InteractiVenn was used to create Venn diagrams (Heberle et al., 2015).

## Results

### Data reproducibility

The repeatability of the Mag-NET technique was assessed using two aliquots of a single sample that were processed in parallel to create technical replicates of the processing workflow, encompassing: enrichment, digestion and LC-MS/MS analysis. Two hundred and twenty-four proteins were included in the analysis, Reduced Major Axis (RMA) regression found a significant (R^2^ = 0.665, P<0.0001) linear relationship between replicates (Supplemental Figure S1A). The 95 % confidence interval (CI) of the intercept and slope were used to assess fixed- and proportional-bias, respectively. The 95 % CI for the Intercept (- 1.4390 to 2.2910) did span zero (i.e. no fixed bias was detected), whereas the 95 % CI for the slope (0.8201– 0.8766) did not span 1, which suggests proportional bias existed between replicate analyses. Coefficient of variation was used to assess the technical variation of protein-specific data The mean CV was 38.1 % (SD 30.6 %) and the median and inter-quartile range in CV was M = 29.7 % (Q1 16.5 % to Q3 53.3 %) (Supplemental Figure S1B).

### EV proteome validation

In total, 449 proteins were confidently identified (>1 unique peptide at FDR <1 %) in plasma EV samples, 411 of which were curated in the Vesiclepedia (Kalra et al., 2012) database and 344 were curated Exocarta (Keerthikumar et al., 2016) database (Figure 2 A & B). Of the 344 proteins that overlapped with Exocarta database, 41 proteins were amongst the top 100 most abundant proteins in exosomes (Figure 2B). After filtering to remove all missing values, 292 proteins were quantified in all (n=24) samples and were subsequently used to investigate statistical differences between Pro, Acd and Xfit groups.

**Figure 2.**
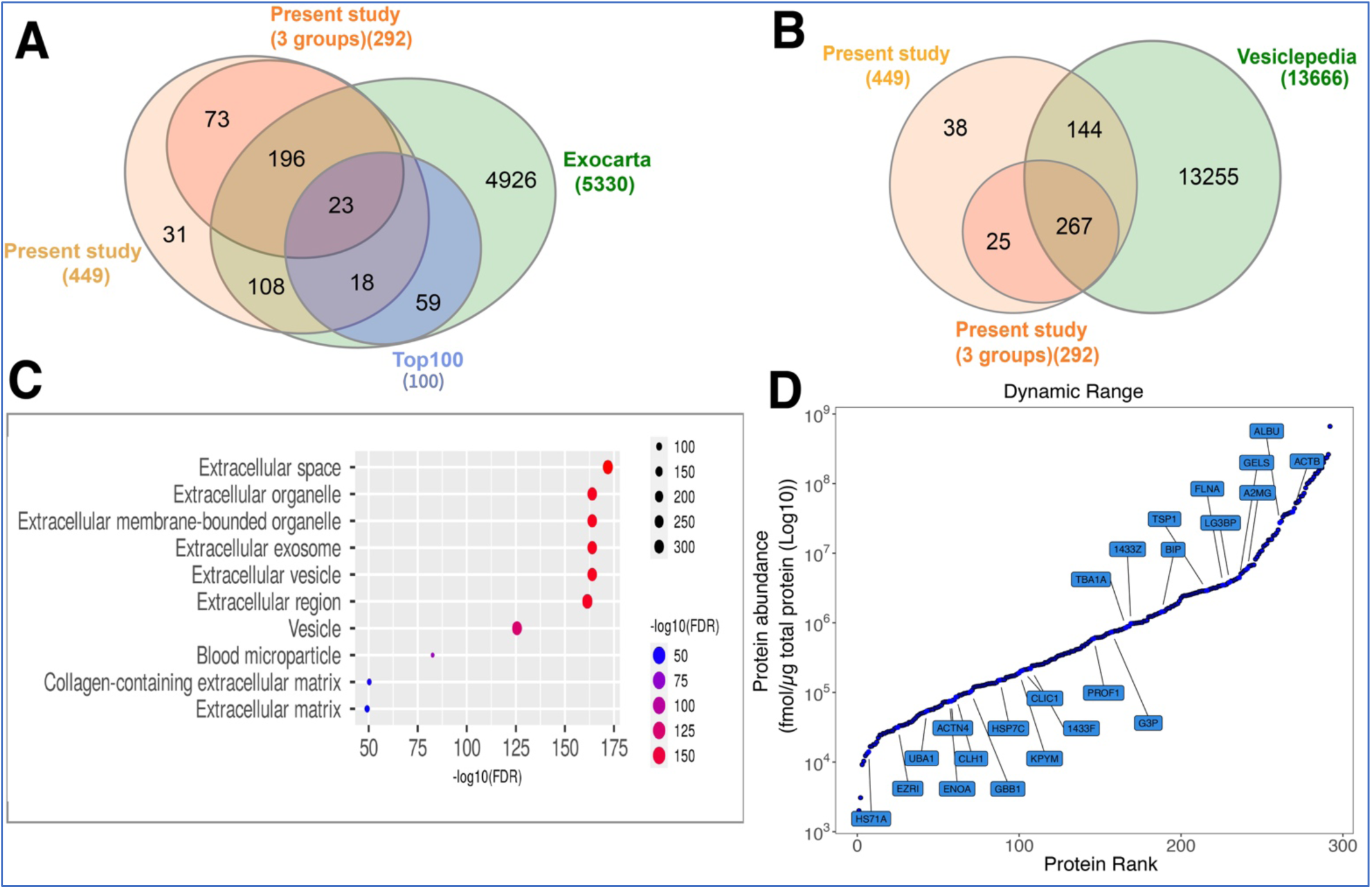
MagResyn enrichment of Plasma EV proteins. Venn diagrams illustrate overlap between proteins in the current study and annotated EV databases: (A) Exocarta and Top 100 exosomes and (B) Vesiclepedia. (C) Gene ontology – Cellular component enrichment analysis of current study against whole human genome. (D) Abundance distribution of 292 proteins ranked (left to right) from lowest to highest abundance. Protein annotations (UniProt accession ids) represent 23 proteins from the top 100 exosomes of the Exocarta database detected in the current study.

The dynamic range of protein abundances included in the statistical analysis spanned from 0.002 to 663.02 nmol/ μg total protein and encompassed 23 proteins from the top 100 exosome proteins reported in Exocarta (Figure 2D). The high abundance proteins included chaperones (heat shock proteins; HS71A, HSP7C, BIP), actin and actin binding proteins (ACTB, FLNA, ACTN4) and proteins of metabolism and transport (ENOA, GBB1, G3P, KPYN, CLIC1) as well as glycoproteins, protein kinase C inhibitors (1433F, 1433Z, TSP1) and ubiquitin related protein (UBA1) (Supplementary Table S1).

One-way analysis of variance (ANOVA) highlighted 128 significant (p < 0.05, FDR <2 %) differences in protein abundance between Acd, Pro and Xfit groups (Figure 3A). Gene ontology cellular components (GO:CC) enriched amongst the significant proteins included Extracellular space [enrichment FDR (eFDR) = 3.50E-149], Extracellular region (eFDR = 2.65E-140), and Extracellular exosomes (eFDR = 7.17E-134) (Figure 2C). Fuzzy c-means clustering segregated the statistically significantly proteins into 5 patterns of abundance difference amongst Acd, Pro and Xfit groups. Protein abundance measurements, statistical outcomes (p- and q-values) and assignments to cluster number are reported in Supplementary Table S2.

**Figure 3.**
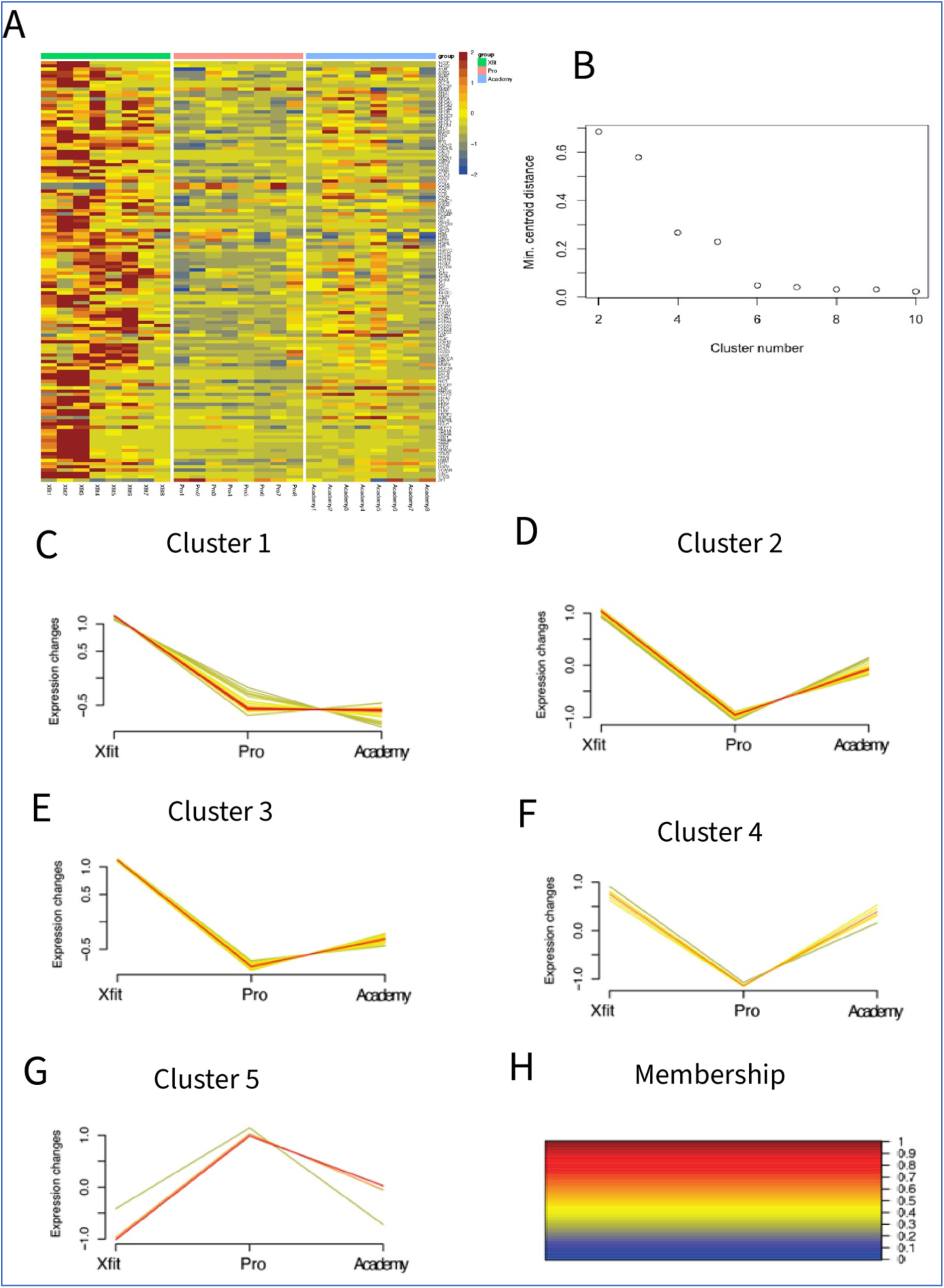
Pattern analysis of protein abundance differences between groups. (A) Heatmap representing the abundance pattern of 128 proteins that exhibited statistically significant differences (One-way ANOVA; p <0.05, q <0.02). (B) Scree plot used to determine the appropriate number of clusters prior to Fuzzy c-means clustering on 128 significantly different proteins. (C-H) Clusters (1 to 5) illustrating patterns of protein abundance differences between Academy, Pro and Xfit athletes with no history in contact sports. (F) Color palette representing the membership value of proteins in each cluster. (Membership reflects the degree of similarity of each protein to the cluster centroid) [Xfit=CrossFit athletes; Pro=Professional rugby players; Academy=Academy rugby players].

Cluster 1 comprised 56 proteins that were specifically more abundant in older athletes without a history of contact sport (Xfit) and were enriched in GO:BP including Platelet aggregation (eFDR = 1.23E-15) and Homotypic cell-cell adhesion (eFDR = 3.14E-14), and KEGG pathways such as Platelet activation (eFDR = 2.74E-08) and ECM-receptor interaction (eFDR = 1.82E-05).

Clusters 2 and 3 encompassed 29 and 28 proteins, respectively, and were most abundant in Xfit athletes, intermediate in Acd players and had the lowest abundance in Pro rugby players. Functional enrichment analysis of Cluster 2 proteins included GO: BP terms involving Complement Activation, classical pathway (eFDR=0.002) and Extracellular matrix organization and structure (eFDR=0.002) and KEGG pathway associated to Complement and coagulation cascades (eFDR=0.0014) and AGE-RAGE signalling pathway in diabetic complications (eFDR=0.027). Functional enrichment analysis of Cluster 3 proteins highlighted enrichment in GO: BP involving Chylomicron remodelling (eFDR = 1.62E-05) and Chylomicron assembly (eFDR = 1.62E-05) and KEGG pathways associated with Cholesterol metabolism (eFDR = 0.0002) and Complement and coagulation cascades (eFDR = 0.0004).

Cluster 4 included 11 proteins that were less abundant in Pro rugby athletes compared to Acd and Xfit groups and was enriched in GO: BP terms, including Low-density lipoprotein particle remodelling (eFDR = 0.0117) and Positive regulation of lipid storage (eFDR = 0.0136). Cluster 5 consisted of four proteins (AMBP, C4A, C4B, SERPINA10), that were more abundant in Pro rugby players only, and were associated with GO: BP terms, positive regulation of apoptotic cell clearance (eFDR = 5.41E-05) and negative regulation of endopeptidase activity (eFDR = 1.25E-06). All Cluster annotations, including enrichment FDR, fold enrichment and GO Pathway enrichment are reported in Supplement Table S2 and Supplement Table S3.

Rugby playing career downregulates proteins involved in Cellular stress response and ECM organization.

Volcano plots were used to illustrate protein-specific detail of the contrasts between pairs of experimental groups (Figure 4). Forty-three proteins exhibited significant difference (P<0.05, q<0.23) between Acd and Pro rugby players. Six proteins (1433F, 1433Z, MYL6,AMBP,ILK, and PON1) were more abundant in Pro players, whereas 37 proteins (OMD, NID1, PCD12, UFO, PI16, FA8, APOA, PEDF, LV140, TM109, CERU, KVD33, LG3BP, BIP, CD44, KV320, HV373, R4RL2, KVD24, MMP9, HV601, CLUS, HV372, LV144, HV5X1, COMP, DSG2, ALS, APOB, LV147, HV118, VCAM1, HGFA, BTD, CD14, LV214, HV323) associated with cell stress responses were less abundant in Pro compared to Acd rugby players (Supplementary Table_S4)

**Figure 4.**
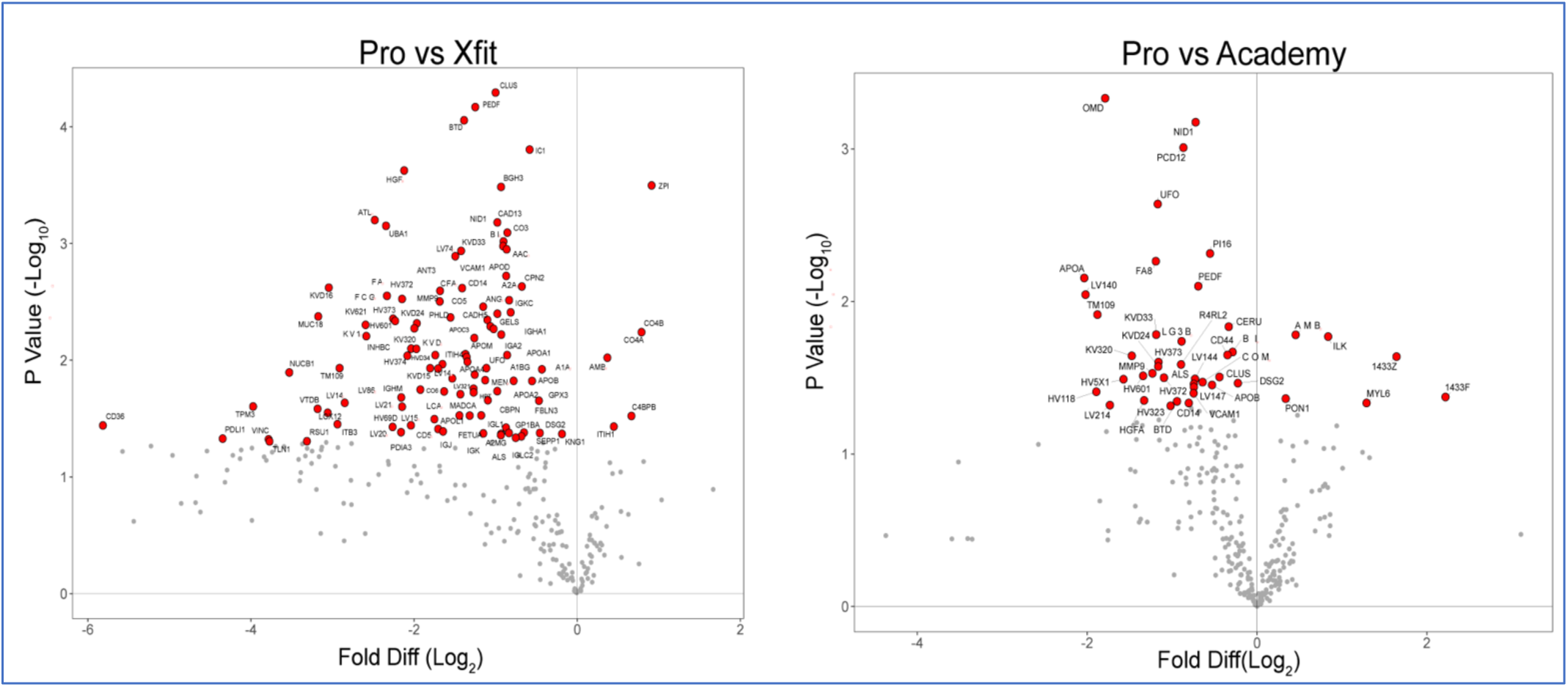
Effect of age and rugby career on protein expression. Visual representation of statistical differences between groups. (A) Volcano plot illustrating log_2_ fold-difference between Pro/Academy, statistically significant differences (P<0.05) are highlighted in red. (B) Volcano plot illustrating log_2_ fold-difference between Pro/ Xfit, statistically significant differences (P<0.05) are highlighted in red. [Xfit=CrossFit athletes; Pro=Professional rugby players; Academy=Academy rugby players; Fold Diff= Fold difference]

Proteins involved in Lipoprotein transport and remodelling are enriched in rugby players.

One-hundred and five proteins were significantly different (one-way ANOVA; P<0.05, q<0.03) between older rugby players and age-matched Xfit athletes with no history of contact sport (Figure 4B; Supplementary Table_S5). Among these, 99 proteins (CLUS, PEDF, BTD, IC1, HGFA, BGH3, ATL4, CAD13, UBA1, CO3, BIP, NID1, AACT, KVD33, LV743, APOD, CPN2, KVD16, VCAM1, ANT3, FA8, HV372, A2AP, CFAH, CD14, IGKC, ANGT, MUC18, PHLD, FCGBP, CO5, HV373, MMP9, KV621, CADH5, KVD24, GELS, IGHA1, KV116, APOC3, HV601, KV320, APOM, IGA2, KVD11, INHBC, ITIH4, UFO, HVD34, A1BG, TM109, HV374, KVD15, A1AT, NUCB1, APOA4, LV147, MENT, APOA1, APOB, LV321, IGHM, APOA2, CO6, HPT, MADCA, LV861, CBPN, GPX3, LV140, TPM3, LV211, VTDB, LOX12, FETUA, APOL1, LCAT, LV151, ITB3, HV69D, CD36, LV208, IGL1, CD5L, IGJ, PDIA3, FBLN3, GP1BA, DSG2, IGK, KNG1, A2MG, ALS, SEPP1, IGLC2, PDLI1, VINC, RSU1, TLN1) were less abundant in samples from Pro players. Six proteins (ZPI, CO4B, CO4A, AMBP, C4BPB, ITIH1) were more abundant in samples from Pro players.

### Analysis of Proteins significantly different only in Pro Rugby players

Figure 5A, illustrates proteins that were significantly different in samples from Pro rugby players compared to either aged-matched athletes with no history of contact sports (Xfit group) or Acd rugby players. Proteins (n=128) that exhibited significant differences (one way ANOVA; p < 0.05; q < 0.02) amongst the 3 experimental groups were plotted based on their fold difference between either Pro rugby players vs age-matched Xfit athletes with no history in contact sports (x-axis) or Pro vs Acd rugby players (y-axis). The majority of proteins occupy the bottom left quadrant and were less abundant in older rugby players than either age-matched Xfit athletes or Academy rugby players. The bottom left quadrant included 30 proteins, which shared GO: BP terms including High-density lipoparticle clearance (eFDR=5.25E-19), Reverse cholesterol transport (eFDR=2.19E-17) and Regulation of plasma lipoprotein oxidation (eFDR=3.01E-17). (Supplementary table S6).

**Figure 5.**
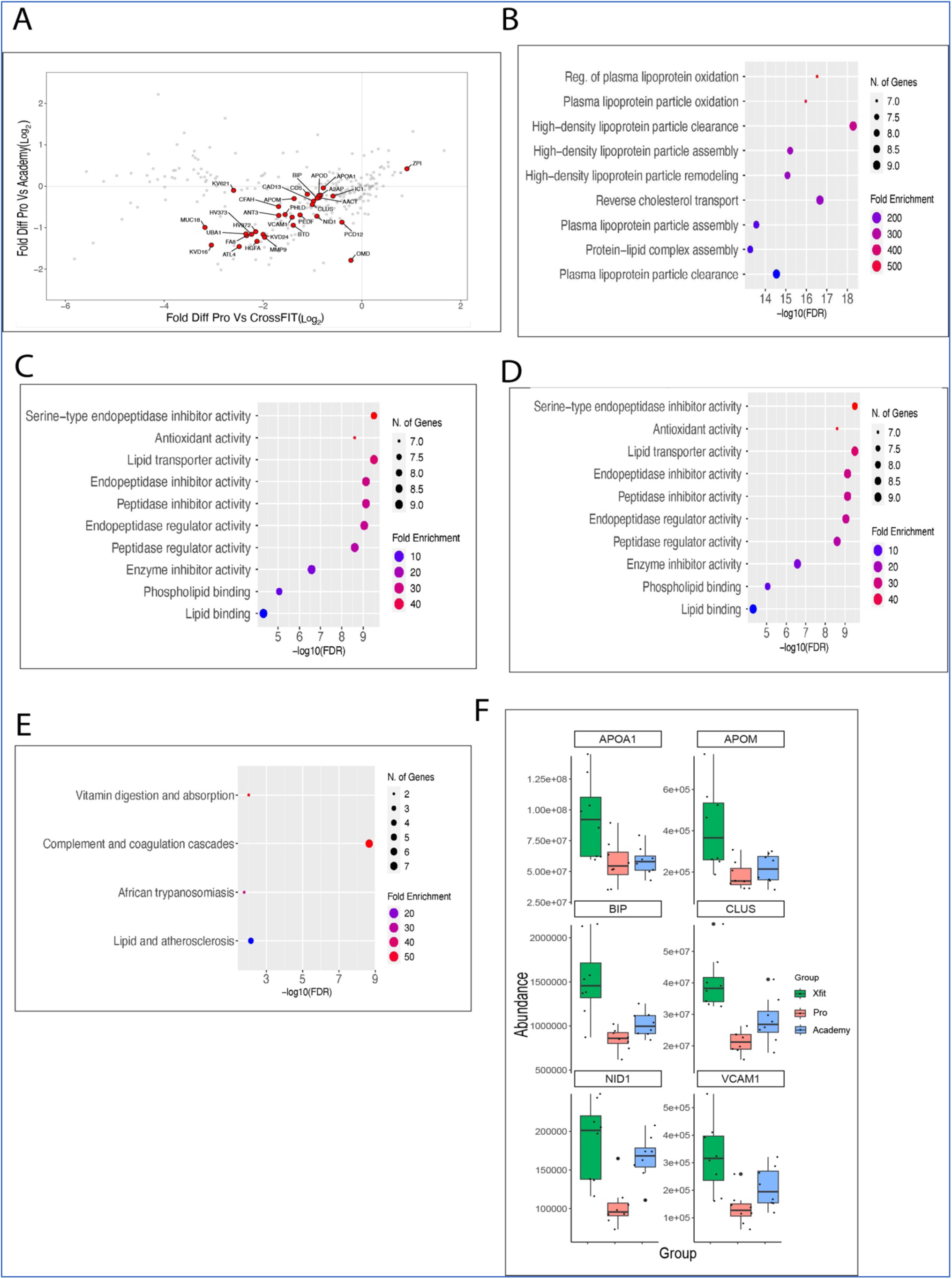
Proteins significantly different in Pro rugby players. Visual representation of statistically significant proteins specific to Pro rugby players. (A) Bottom left quadrant of a scatterplot comparing protein abundance log_2_ fold-difference between Pro/ Xfit (x-axis) and Pro/ Academy (y-axis). Statistically significant (P<0.05, FDR <2 %) proteins are coloured red and annotated with UniProt identifiers. (B to E) Functional enrichment analysis of 21 proteins from lower left quadrant of scatterplot, including Biological process (B), Cellular component (C), Molecular Function (D) and KEGG pathway (E). (F) Box plots of 6 proteins belong to the top-ranked biological process, “cellular response to reactive oxygen species.” [Xfit=CrossFit athletes; Pro=Professional rugby players; Academy=Academy rugby players; Fold Diff. = Fold difference]

## Discussion

The risk of neurological damage to competitors in contact sports including rugby is a serious concern and currently little is known about the mechanisms that may predispose athletes with long careers in contact sports to a greater risk of neurological disease. Plasma biomarkers offer an opportunity to objectively assess biological differences and, in particular, non-targeted analysis of the EV proteome may generate new insight into processes involved in neurological disorders. Seven of the proteins (APOA1, APOM, CLUS, BIP, VCAM1, NID1 and MMP9) that were depleted and one protein ZPI that was elevated in EV samples from Pro rugby players have previously recognised roles in neurodegenerative processes. Lower levels of each of these proteins have been associated with various neurological conditions, and their shared response in Pro rugby players could be the basis of a biomarker panel for monitoring the accumulative consequences of sports injuries.

Low levels of apolipoprotein A-I (APOA1) correlate with greater risk of ischemic strokes and neurodegeneration, and lower serum APOA1 levels may contribute to metabolic dysregulation and neuroinflammation following concussions (Ohtani et al., 2020). Similarly, apolipoprotein M (APOM), which plays a crucial role in the anti-inflammatory properties of high-density lipoprotein (HDL) (Christoffersen et al., 2011), has been linked to impaired endothelial function and increased inflammation when APOM levels are reduced (Christensen et al., 2016). Clusterin (CLUS), also known as apolipoprotein J (APOJ), is an extracellular chaperone that is strongly associated with AD (Foster et al., 2019). CLUS promotes the clearance of toxic amyloid-beta (Aβ) peptides (Wojtas et al., 2020), and variants in the CLUS gene are an acknowledged marker of late onset AD (Szymanski et al., 2011). CLUS also has neuroprotective roles during acute events such as perioperative ischemic-reperfusion injury (Iłżecka et al., 2019), which suggests CLUS deficiency could exacerbate neuronal damage following concussive sports injuries.

The endoplasmic reticulum chaperone (BiP), also known as GRP78, is critical for intracellular stress responses, and the endoplasmic reticulum is also intimately connected with EV formation (Ye and Liu, 2022). BiP may be particularly relevant during recovery from sports-related concussions when the brain experiences metabolic disturbances and increases in oxidative stress. BiP expression is increased in experimental models of traumatic brain injury (Truettner et al., 2007), whereas lower levels of BiP are associated with greater susceptibility to cellular stress and may contribute to conditions such as AD (Kumar et al., 2022). BiP is also exposed on the outer membrane of brain microvascular endothelia cells and autoantibodies to GRP78 modulate blood brain barrier (BBB) function (Shimizu et al., 2017). Similarly, vascular cell adhesion molecule 1 (VCAM1) is an adhesion molecule protein expressed on endothelial cells and is involved in maintaining BBB integrity. In the chronic cerebral hypoperfusion model of vascular dementia, VCAM1 expression is increased in brain endothelia cells and correlates with BBB dysfunction (Zhang, 2024). Plasma levels of VCAM1 correlate strongly with age in humans and mice, and brain endothelia cells exposed to plasma from aged mice upregulate VCAM1 abundance (Yousef et al., 2019). This suggests low circulating levels of VCAM1 may be beneficial, potentially limiting excessive inflammation and maintaining BBB integrity, which is critical for neuronal health.

The vascular basement membrane is a component of the BBB (Thomsen et al., 2017) and 2 proteins, Nidogen-1 (NID1) and matrix metalloproteinase 9 (MMP9), that were depleted in samples from Pro rugby players are associated with the basement membrane. NID1 is a key structural component of the basement membrane (BM) that interlinks with and stabilises laminin and collagen IV, and is crucial for neuronal health and function (Vasudevan et al., 2010). NID1 interacts with reelin which is a basement membrane protein that was more abundant in plasma EV of former NFL players (Muraoka et al., 2021). Matrix metalloproteinase 9 (MMP9), is also implicated in the stabilization and maintenance of the extracellular matrix (ECM) and patients with AD exhibit lower levels of plasma MMP9 (Gong and Jia, 2022).

Recently, plasma EV have emerged as a cascading multisystem physiological adaptation that trigger immune responses (Chong et al., 2024). Circulating EV have also been shown to exhibit anti-inflammatory effects in response to varied training regimes (Maggio et al., 2023) and exercise-induced muscle damage is associated with changes to plasma EV profile (Kyriakidou et al., 2021). One protein, Z-dependent protease inhibitor (ZPI), also known as SERPINA10, was elevated in Pro samples. ZPI is known to form a complex with Protein Z (PZ), which enhances the ability of PZ to inhibit coagulation factors, including factor Xa and factor Xia (Han et al., 1999). The actions of ZPI prevent excessive thrombin generation during the early stages of coagulation (Zhang et al., 2008) and reduce thrombotic risk. ZPI may also have protective roles in endothelial injury by preventing oxidized low-density lipoprotein (ox-LDL)-mediated endothelial injury and inhibiting processes, including endothelial-to-mesenchymal transition, inflammation, and apoptosis (Zhang et al., 2022).

In this study, we employed the Mag-Net method reported (Wu et al., 2023) that relies on both size-based ‘netting’ and charge-based binding to enrich membrane bound particles, including extracellular vesicles (EV) from blood plasma samples. We compared our dataset with manually curated EV databases (Exocarta and Vesiclepedia) and confirmed the selective enrichment of EV proteins. Nevertheless, a good proportion of the putative biomarkers highlighted in our analysis of Pro rugby samples were apolipoproteins that are more likely associated with lipid particles such as HDL. Lipid particles are more abundant in plasma than EV and the protein composition of lipid particles and EV differ. Apolipoprotein APOA1, which is a marker of HDL and chylomicron particles, is co-extracted by traditional EV isolation workflows that rely on size exclusion chromatography (Karimi et al., 2018). Wu reports Mag-Net depletes lipoproteins are depleted ∼80 % compared to raw plasma and, specifically, APOA1 was -2.5 (log2) less in enriched samples (Wu et al., 2023) but the apolipoprotein biomarkers highlighted in the current work are unlikely to be of EV origin.

Because EV are able to traverse the blood-brain barrier, research on SRC has explored plasma EV as a potential source of biomarkers (Meier et al., 2022). However, it is important to clarify our study did not collect data on the incidence or type of injuries experienced by our participants. We also cannot rule out the contribution of musculoskeletal injuries, which occur frequently in Rugby, or lifestyle factors that could affect the plasma EV profile. Future studies are required to verify whether the biomarker pattern discovered herein persists when physical activity level, recent injury history and sample collection protocols are controlled. Nevertheless, despite samples from Acd and Pro players being collected in different pre-season phases and the lack of control over the time between sample collection and the last bout of exercise along with limited dietary control, we were able to detect consistent differences in the plasma EV proteome.

## Conclusion

Non-targeted analysis highlighted that proteins associated with neuroprotection were specifically depleted in the plasma EV proteome of long-serving professional rugby players compared to younger academy rugby players or age-matched cross-fit athletes that did not have a history of collision-related sports injuries. Our findings bring new insight into the biological processes affected by a professional rugby playing career but further studies are required to test whether the putative biomarkers reported herein are linked to sports-related concussion and musculoskeletal injuries.

## References

McCrory, P.; Meeuwisse, W.; Dvorak, J.; Aubry, M.; Bailes, J.; Broglio, S.; Cantu, R.C.; Cassidy, D.; Echemendia, R.J.; Castellani, R.J. Consensus statement on concussion in sport—the 5th international conference on concussion in sport held in Berlin, October 2016. British journal of sports medicine. 2017;51(11):838-847.

PRISP. England Professional Rugby Injury Surveillance Project 2022–2023 Season Report. 2024.

Broglio, S.P.; McAllister, T.; Katz, B.P.; LaPradd, M.; Zhou, W.; McCrea, M.A.; Hoy, A.; Hazzard, J.B.; Kelly, L.A.; DiFiori, J.; Ortega, J.D.; Port, N.; Putukian, M.; Langford, D.; McDevitt, J.; Campbell, D.; Jackson, J.C.; McGinty, G.; Estevez, C.; Cameron, K.L.; Houston, M.N.; Svoboda, S.J.; Susmarski, A.J.; Giza, C.; Benjamin, H.J.; Kaminski, T.W.; Buckley, T.; Clugston, J.R.; Schmidt, J.; Feigenbaum, L.A.; Eckner, J.T.; Mihalik, J.; Miles, J.D.; Anderson, S.; Arbogast, K.; Master, C.L.; Kontos, A.P.; Chrisman, S.P.D.; Brooks, M.A.; Rowson, S.; Duma, S.M.; Miles, C.; Investigators, C.C. The Natural History of Sport-Related Concussion in Collegiate Athletes: Findings from the NCAA-DoD CARE Consortium. Sports Medicine. 2022 2022/02/01;52(2):403-415.

Blecher, R.; Elliott, M.A.; Yilmaz, E.; Dettori, J.R.; Oskouian, R.J.; Patel, A.; Clarke, A.; Hutton, M.; McGuire, R.; Dunn, R.; DeVine, J.; Twaddle, B.; Chapman, J.R. Contact Sports as a Risk Factor for Amyotrophic Lateral Sclerosis: A Systematic Review. Global Spine Journal. 2019;9(1):104–118.

Chen, G.X.; Douwes, J.; van den Berg, L.H.; Glass, B.; McLean, D.; ’t Mannetje, A.M. Sports and trauma as risk factors for Motor Neurone Disease: New Zealand case–control study. Acta Neurologica Scandinavica. 2022;145(6):770–785.

Hume, P.A.; Theadom, A.; Lewis, G.N.; Quarrie, K.L.; Brown, S.R.; Hill, R.; Marshall, S.W. A comparison of cognitive function in former rugby union players compared with former non-contact-sport players and the impact of concussion history. Sports medicine. 2017;47:1209–1220.

Gallo, V.; McElvenny, D.M.; Seghezzo, G.; Kemp, S.; Williamson, E.; Lu, K.; Mian, S.; James, L.; Hobbs, C.; Davoren, D.; Arden, N.; Davies, M.; Malaspina, A.; Loosemore, M.; Stokes, K.; Cross, M.; Crutch, S.; Zetterberg, H.; Pearce, N. Concussion and long-term cognitive function among rugby players—The BRAIN Study. Alzheimer’s & Dementia. 2022;18(6):1164–1176.

Gosselin, N.; Bottari, C.; Chen, J.-K.; Huntgeburth, S.C.; De Beaumont, L.; Petrides, M.; Cheung, B.; Ptito, A. Evaluating the cognitive consequences of mild traumatic brain injury and concussion by using electrophysiology. Neurosurgical focus. 2012;33(6):E7.

Chen, J.-K.; Johnston, K.M.; Petrides, M.; Ptito, A. Recovery from mild head injury in sports: evidence from serial functional magnetic resonance imaging studies in male athletes. Clinical Journal of Sport Medicine. 2008;18(3):241–247.

Bartnik-Olson, B.L.; Holshouser, B.; Wang, H.; Grube, M.; Tong, K.; Wong, V.; Ashwal, S. Impaired neurovascular unit function contributes to persistent symptoms after concussion: a pilot study. Journal of neurotrauma. 2014;31(17):1497–1506.

Shahim, P.; Tegner, Y.; Marklund, N.; Blennow, K.; Zetterberg, H. Neurofilament light and tau as blood biomarkers for sports-related concussion. Neurology. 2018;90(20):e1780–e1788.

Laverse, E.; Guo, T.; Zimmerman, K.; Foiani, M.S.; Velani, B.; Morrow, P.; Adejuwon, A.; Bamford, R.; Underwood, N.; George, J.; Brooke, D.; O’Brien, K.; Cross, M.J.; Kemp, S.P.T.; Heslegrave, A.J.; Hardy, J.; Sharp, D.J.; Zetterberg, H.; Morris, H.R. Plasma glial fibrillary acidic protein and neurofilament light chain, but not tau, are biomarkers of sports-related mild traumatic brain injury. Brain Communications. 2020;2(2).

Oris, C.; Durif, J.; Rouzaire, M.; Pereira, B.; Bouvier, D.; Kahouadji, S.; Abbot, M.; Brailova, M.; Lehmann, S.; Hirtz, C.; Decq, P.; Dusfour, B.; Marchi, N.; Sapin, V. Blood Biomarkers for Return to Play after Concussion in Professional Rugby Players. J Neurotrauma. 2023 Feb;40(3-4):283–295.

Meier, T.B.; Nelson, L.D.; Huber, D.L.; Bazarian, J.J.; Hayes, R.L.; McCrea, M.A. Prospective Assessment of Acute Blood Markers of Brain Injury in Sport-Related Concussion. J Neurotrauma. 2017 Nov 15;34(22):3134–3142.

Stocchero, C.M.; Oses, J.P.; Cunha, G.S.; Martins, J.B.; Brum, L.M.; Zimmer, E.R.; Souza, D.O.; Portela, L.V.; Reischak-Oliveira, A. Serum S100B level increases after running but not cycling exercise. Appl Physiol Nutr Metab. 2014 Mar;39(3):340–344.

Zhao, J.; Wang, Y.; Zhao, D.; Zhang, L.; Chen, P.; Xu, X. Integration of metabolomics and proteomics to reveal the metabolic characteristics of high-intensity interval training. Analyst. 2020;145(20):6500–6510.

Mi, M.Y.; Barber, J.L.; Rao, P.; Farrell, L.A.; Sarzynski, M.A.; Bouchard, C.; Robbins, J.M.; Gerszten, R.E. Plasma Proteomic Kinetics in Response to Acute Exercise. Molecular & Cellular Proteomics. 2023;22(8).

Meier, T.B.; Guedes, V.A.; Smith, E.G.; Sass, D.; Mithani, S.; Vorn, R.; Savitz, J.; Teague, T.K.; McCrea, M.A.; Gill, J.M. Extracellular vesicle-associated cytokines in sport-related concussion. Brain, Behavior, and Immunity. 2022 2022/02/01/;100:83-87.

Rath, M.; Sayoc, J.; Burns, K.; McDevitt, J.; Fan, X.; Tierney, R.; Wu, J.; Park, J.Y. Glial cell-Derived, but Not Neuron-Derived, Extracellular Vesicles May Serve as Novel Biomarkers of Acute Sport-Related Concussion. The FASEB Journal. 2022;36(S1).

Frühbeis, C.; Helmig, S.; Tug, S.; Simon, P.; Krämer-Albers, E.-M. Physical exercise induces rapid release of small extracellular vesicles into the circulation. Journal of Extracellular Vesicles. 2015;4(1):28239.

Zaborowski, M.P.; Balaj, L.; Breakefield, X.O.; Lai, C.P. Extracellular Vesicles: Composition, Biological Relevance, and Methods of Study. BioScience. 2015;65(8):783–797.

Banks, W.A.; Sharma, P.; Bullock, K.M.; Hansen, K.M.; Ludwig, N.; Whiteside, T.L. Transport of Extracellular Vesicles across the Blood-Brain Barrier: Brain Pharmacokinetics and Effects of Inflammation. International Journal of Molecular Sciences. 2020;21(12):4407.

Osteikoetxea, X.; Sódar, B.; Németh, A.; Szabó-Taylor, K.; Pálóczi, K.; Vukman, K.V.; Tamási, V.; Balogh, A.; Kittel, Á.; Pállinger, É.; Buzás, E.I. Differential detergent sensitivity of extracellular vesicle subpopulations11Electronic supplementary information (ESI) available. See DOI: 10.1039/c5ob01451d. Organic & Biomolecular Chemistry. 2015 2015/10/01/;13(38):9775-9782.

Ko, J.; Hemphill, M.; Yang, Z.; Sewell, E.; Na, Y.J.; Sandsmark, D.K.; Haber, M.; Fisher, S.A.; Torre, E.A.; Svane, K.C.; Omelchenko, A.; Firestein, B.L.; Diaz-Arrastia, R.; Kim, J.; Meaney, D.F.; Issadore, D. Diagnosis of traumatic brain injury using miRNA signatures in nanomagnetically isolated brain-derived extracellular vesicles. Lab Chip. 2018 Dec 7;18(23):3617–3630.

Muraoka, S.; DeLeo, A.M.; Yang, Z.; Tatebe, H.; Yukawa-Takamatsu, K.; Ikezu, S.; Tokuda, T.; Issadore, D.; Stern, R.A.; Ikezu, T. Proteomic Profiling of Extracellular Vesicles Separated from Plasma of Former National Football League Players at Risk for Chronic Traumatic Encephalopathy. Aging Dis. 2021 Sep;12(6):1363–1375.

Brahmer, A.; Neuberger, E.; Esch-Heisser, L.; Haller, N.; Jorgensen, M.M.; Baek, R.; Möbius, W.; Simon, P.; Krämer-Albers, E.-M. Platelets, endothelial cells and leukocytes contribute to the exercise-triggered release of extracellular vesicles into the circulation. Journal of Extracellular Vesicles. 2019;8(1):1615820.

Holcar, M.; Ferdin, J.; Sitar, S.; Tušek-Žnidarič, M.; Dolžan, V.; Plemenitaš, A.; Žagar, E.; Lenassi, M. Enrichment of plasma extracellular vesicles for reliable quantification of their size and concentration for biomarker discovery. Scientific Reports. 2020 2020/12/07;10(1):21346.

Wu, C.C.; Tsantilas, K.A.; Park, J.; Plubell, D.; Naicker, P.; Govender, I.; Buthelezi, S.; Stoychev, S.; Jordaan, J.; Merrihew, G.; Huang, E.; Parker, E.D.; Riffle, M.; Hoofnagle, A.N.; MacCoss, M.J. Mag-Net: Rapid enrichment of membrane-bound particles enables high coverage quantitative analysis of the plasma proteome. 2023.

Hesketh, S.J.; Stansfield, B.N.; Stead, C.A.; Burniston, J.G. The application of proteomics in muscle exercise physiology. Expert Rev Proteomics. 2020 Nov-Dec;17(11-12):813-825.

Burniston, J.G.; Connolly, J.; Kainulainen, H.; Britton, S.L.; Koch, L.G. Label-free profiling of skeletal muscle using high-definition mass spectrometry. PROTEOMICS. 2014;14(20):2339–2344.

Ge, S.X.; Jung, D.; Yao, R. ShinyGO: a graphical gene-set enrichment tool for animals and plants. Bioinformatics. 2019;36(8):2628–2629.

Heberle, H.; Meirelles, G.V.; da Silva, F.R.; Telles, G.P.; Minghim, R. InteractiVenn: a web-based tool for the analysis of sets through Venn diagrams. BMC Bioinformatics. 2015 2015/05/22;16(1):169.

Kalra, H.; Simpson, R.J.; Ji, H.; Aikawa, E.; Altevogt, P.; Askenase, P.; Bond, V.C.; Borràs, F.E.; Breakefield, X.; Budnik, V.; Buzas, E.; Camussi, G.; Clayton, A.; Cocucci, E.; Falcon-Perez, J.M.; Gabrielsson, S.; Gho, Y.S.; Gupta, D.; Harsha, H.C.; Hendrix, A.; Hill, A.F.; Inal, J.M.; Jenster, G.; Krämer-Albers, E.-M.; Lim, S.K.; Llorente, A.; Lötvall, J.; Marcilla, A.; Mincheva-Nilsson, L.; Nazarenko, I.; Nieuwland, R.; Nolte-’t Hoen, E.N.M.; Pandey, A.; Patel, T.; Piper, M.G.; Pluchino, S.; Prasad, T.S.K.; Rajendran, L.; Raposo, G.; Record, M.; Reid, G.E.; Sánchez-Madrid, F.; Schiffelers, R.M.; Siljander, P.; Stensballe, A.; Stoorvogel, W.; Taylor, D.; Thery, C.; Valadi, H.; van Balkom, B.W.M.; Vázquez, J.; Vidal, M.; Wauben, M.H.M.; Yáñez-Mó, M.; Zoeller, M.; Mathivanan, S. Vesiclepedia: A Compendium for Extracellular Vesicles with Continuous Community Annotation. PLOS Biology. 2012;10(12):e1001450.

Keerthikumar, S.; Chisanga, D.; Ariyaratne, D.; Al Saffar, H.; Anand, S.; Zhao, K.; Samuel, M.; Pathan, M.; Jois, M.; Chilamkurti, N.; Gangoda, L.; Mathivanan, S. ExoCarta: A Web-Based Compendium of Exosomal Cargo. Journal of Molecular Biology. 2016 2016/02/22/;428(4):688-692.

Ohtani, R.; Nirengi, S.; Sakane, N. Association Between Serum Apolipoprotein A1 Levels, Ischemic Stroke Subtypes and Plaque Properties of the Carotid Artery. Journal of Clinical Medicine Research. 2020;12(9):598–603.

Christoffersen, C.; Obinata, H.; Kumaraswamy, S.B.; Galvani, S.; Ahnström, J.; Sevvana, M.; Egerer-Sieber, C.; Muller, Y.A.; Hla, T.; Nielsen, L.B.; Dahlbäck, B. Endothelium-Protective Sphingosine-1-Phosphate Provided by HDL-associated Apolipoprotein M. Proceedings of the National Academy of Sciences. 2011;108(23):9613–9618.

Christensen, P.H.; Liu, C.H.; Swendeman, S.; Obinata, H.; Qvortrup, K.; Nielsen, L.B.; Hla, T.; Lorenzo, A.D.; Christoffersen, C. Impaired Endothelial Barrier Function in Apolipoprotein M-deficient Mice Is Dependent on Sphingosine-1-phosphate Receptor 1. The Faseb Journal. 2016;30(6):2351–2359.

Foster, E.M.; Dangla-Valls, A.; Lovestone, S.; Ribe, E.M.; Buckley, N.J. Clusterin in Alzheimer’s Disease: Mechanisms, Genetics, and Lessons From Other Pathologies. Front Neurosci. 2019;13:164.

Wojtas, A.; Sens, J.P.; Kang, S.S.; Baker, K.E.; Berry, T.J.; Kurti, A.; Daughrity, L.M.; Jansen-West, K.; Dickson, D.W.; Petrucelli, L.; Bu, G.; Liu, C.-C.; Fryer, J.D. Astrocyte-Derived Clusterin Suppresses Amyloid Formation in Vivo. Molecular Neurodegeneration. 2020;15(1).

Szymanski, M.; Wang, R.; Bassett, S.S.; Avramopoulos, D. Alzheimer’s risk variants in the clusterin gene are associated with alternative splicing. Transl Psychiatry. 2011;1(7):e18-.

Iłżecka, J.; Iłżecki, M.; Grabarska, A.; Dave, S.; Feldo, M.; Zubilewicz, T. Clusterin as a Potential Marker of Brain Ischemia-Reperfusion Injury in Patients Undergoing Carotid Endarterectomy. Upsala Journal of Medical Sciences. 2019;124(3):193–198.

Ye, J.; Liu, X. Interactions between endoplasmic reticulum stress and extracellular vesicles in multiple diseases. Frontiers in Immunology. 2022 2022-August-11;13.

Truettner, J.S.; Hu, B.; Alonso, O.F.; Bramlett, H.M.; Kokame, K.; Dietrich, W.D. Subcellular Stress Response after Traumatic Brain Injury. Journal of Neurotrauma. 2007;24(4):599–612.

Kumar, V.; Kim, S.H.; Bishayee, K. Dysfunctional Glucose Metabolism in Alzheimer’s Disease Onset and Potential Pharmacological Interventions. Int J Mol Sci. 2022 Aug 23;23(17).

Shimizu, F.; Schaller, K.L.; Owens, G.P.; Cotleur, A.C.; Kellner, D.; Takeshita, Y.; Obermeier, B.; Kryzer, T.J.; Sano, Y.; Kanda, T.; Lennon, V.A.; Ransohoff, R.M.; Bennett, J.L. Glucose-regulated protein 78 autoantibody associates with blood-brain barrier disruption in neuromyelitis optica. Sci Transl Med. 2017 Jul 5;9(397).

Zhang, H. Increased Expression of VCAM1 on Brain Endothelial Cells Drives Blood–Brain Barrier Impairment Following Chronic Cerebral Hypoperfusion. Acs Chemical Neuroscience. 2024;15(10):2028–2041.

Yousef, H.; Czupalla, C.J.; Lee, D.; Chen, M.B.; Burke, A.N.; Zera, K.A.; Zandstra, J.; Berber, E.; Lehallier, B.; Mathur, V.; Nair, R.V.; Bonanno, L.; Yang, A.; Peterson, T.C.; Hadeiba, H.; Merkel, T.; Körbelin, J.; Schwaninger, M.; Buckwalter, M.S.; Quake, S.R.; Butcher, E.C.; Wyss-Coray, T. Aged Blood Impairs Hippocampal Neural Precursor Activity and Activates Microglia via Brain Endothelial Cell VCAM1. Nature Medicine. 2019;25(6):988–1000.

Thomsen, M.S.; Routhe, L.J.; Moos, T. The vascular basement membrane in the healthy and pathological brain. J Cereb Blood Flow Metab. 2017 Oct;37(10):3300–3317.

Vasudevan, A.; Ho, M.S.; Weiergräber, M.; Nischt, R.; Schneider, T.; Lie, A.; Smyth, N.; Köhling, R. Basement membrane protein nidogen-1 shapes hippocampal synaptic plasticity and excitability. Hippocampus. 2010;20(5):608–620.

Gong, M.; Jia, J. Contribution of Blood-Brain Barrier-Related Blood-Borne Factors for Alzheimer’s Disease vs. Vascular Dementia Diagnosis: A Pilot Study. Frontiers in Neuroscience. 2022;16.

Chong, M.C.; Shah, A.D.; Schittenhelm, R.B.; Silva, A.; James, P.F.; Wu, S.S.X.; Howitt, J. Acute exercise-induced release of innate immune proteins via small extracellular vesicles changes with aerobic fitness and age. Acta Physiologica. 2024;240(3):e14095.

Maggio, S.; Canonico, B.; Ceccaroli, P.; Polidori, E.; Cioccoloni, A.; Giacomelli, L.; Ferri Marini, C.; Annibalini, G.; Gervasi, M.; Benelli, P.; Fabbri, F.; Del Coco, L.; Fanizzi, F.P.; Giudetti, A.M.; Lucertini, F.; Guescini, M. Modulation of the Circulating Extracellular Vesicles in Response to Different Exercise Regimens and Study of Their Inflammatory Effects. International Journal of Molecular Sciences. 2023;24(3):3039.

Kyriakidou, Y.; Cooper, I.; Kraev, I.; Lange, S.; Elliott, B.T. Preliminary Investigations Into the Effect of Exercise-Induced Muscle Damage on Systemic Extracellular Vesicle Release in Trained Younger and Older Men. Front Physiol. 2021;12:723931.

Han, X.; Huang, Z.F.; Fiehler, R.; Broze, G.J., Jr. The protein Z-dependent protease inhibitor is a serpin. Biochemistry. 1999 Aug 24;38(34):11073–11078.

Zhang, J.; Tu, Y.; Lu, L.; Lasky, N.; Broze, G.J., Jr. Protein Z-dependent protease inhibitor deficiency produces a more severe murine phenotype than protein Z deficiency. Blood. 2008 May 15;111(10):4973–4978.

Zhang, X.; Li, Z.; Liu, X.; Qin, X.; Luo, J.; Zhang, W.; Liu, B.; Wei, Y. ZPI prevents ox-LDL-mediated endothelial injury leading to inhibition of EndMT, inflammation, apoptosis, and oxidative stress through activating Pi3k/Akt signal pathway. Drug Dev Res. 2022 Aug;83(5):1212–1225.

Karimi, N.; Cvjetkovic, A.; Jang, S.C.; Crescitelli, R.; Hosseinpour Feizi, M.A.; Nieuwland, R.; Lotvall, J.; Lasser, C. Detailed analysis of the plasma extracellular vesicle proteome after separation from lipoproteins. Cell Mol Life Sci. 2018 Aug;75(15):2873–2886.

