## Supplement Figure 1 for "Non-targeted Analysis of Extracellular Vesicle-Enriched Plasma Proteome between Early and Late Rugby Playing Career"

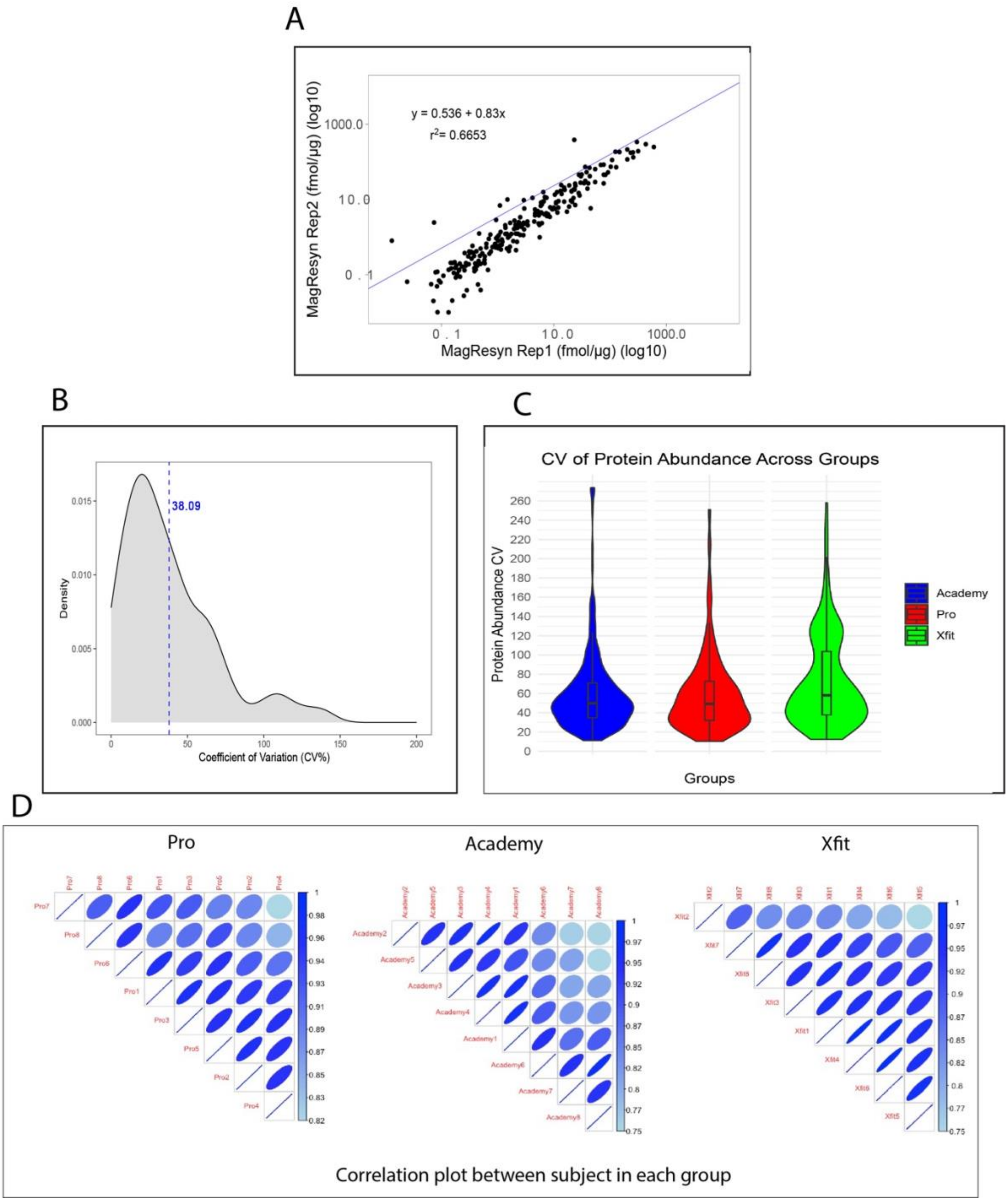


Supplement Figure1: (A)Repeatability (RMA)analysis between two replicates of same sample using MagResyn. The black dots represent protein abundance of each protein quantified across both replicates. (B)Denisty plot represnting Cumulative variation(CV) Coefficient of variation was used to assess variability of protein-specific data (C)CV of protein abudance Across groups.(D)Correlation plot showing correlation (r values) of proteins abundance between subjects in the same group.
